# MetaGIN: A Lightweight Framework for Molecular Property Prediction

**DOI:** 10.1101/2023.10.18.562883

**Authors:** Xuan Zhang, Cheng Chen, Xiaoting Wang, Haitao Jiang, Wei Zhao, Xuefeng Cui

## Abstract

Recent advancements in AI-based synthesis of small molecules have led to the creation of extensive databases, housing billions of small molecules. Given this vast scale, traditional quantum chemistry (QC) methods become inefficient for determining the chemical and physical properties of such an extensive array of molecules. As these properties are key to new drug research and development, deep learning techniques have recently been developed. Here, we present MetaGIN, a lightweight framework designed for molecular property prediction.

While traditional GNN models with 1-hop edges (i.e., covalent bonds) are sufficient for abstract graph representation, they are inadequate for capturing 3D features. Our MetaGIN model shows that including 2-hop and 3-hop edges (representing bond and torsion angles, respectively) is crucial to fully comprehend the intricacies of 3D molecules. Moreover, MetaGIN is a streamlined model with fewer than 10 million parameters, making it ideal for fine-tuning on a single GPU. It also adopts the widely acknowledged MetaFormer framework, which has consistently shown high accuracy in many computer vision tasks.

Through our experiments, MetaGIN achieves a mean absolute error (MAE) of 0.0851 with just 8.87M parameters on the PCQM4Mv2 dataset. Furthermore, MetaGIN outperforms leading techniques, showcasing superior performance across several datasets in the MoleculeNet benchmark.

## 1 Introduction

Molecular property prediction, as a task in quantum chemistry (QC), plays the vital role in drug discovery [1]. The properties of molecules assist scientists in screening more benign and effective molecules from thousands of drug candidates [2,3]. Over the past few decades, several QC methods for obtaining these properties have been proposed, including density functional theory (DFT) [4], quantum Monte Carlo methods [5], and coupled cluster (CC) approaches [6]. However, these methods are both time-consuming and computationally demanding. For instance, DFT, the most widely used method in QC, can take anywhere from hours to days to calculate the properties of a single molecule. Given the rapid development of large-scale molecular databases, traditional methods have become impractical for calculating the properties of all molecules in these databases.

With the advancement of deep learning, a new avenue has opened up where artificial intelligence can assist in molecular property prediction [7, 8] . For example, several models have demonstrated impressive performance on the PCQM4Mv2 dataset [9]. This performance is depicted in Fig. 1, which focuses on models that do not utilize 3D structure data. Among these, Transformer-based architectures [10], such as GEM-2 [11], GPS [12], EGT [13], and GRPE [14], frequently outperform alternative methods. However, these Transformer-like architectures often have extensive depth and breadth, leading to a high number of parameters and significant hardware requirements. Such complexity creates challenges, especially for researchers and organizations with limited hardware resources, complicating both the training and fine-tuning stages.

**Fig. 1.**
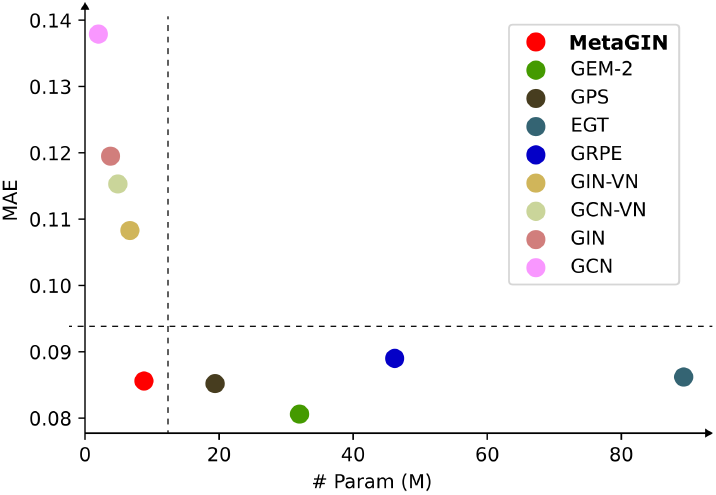
Performance evaluation on the PCQM4Mv2 dataset (without 3D structures). MetaGIN achieves a nearly optimal Mean Absolute Error (MAE) with a compact model that contains fewer than 10 million parameters.

On the other hand, graph-based neural networks (GNNs) generally have fewer parameters and perform inference more quickly than Transformer-like models. As illustrated in Fig. 1, graph-based models, such as GCN [15], GIN [16] and their variants, are highlighted in lighter colors and positioned in the upper-left quadrant. This suggests that these models usually have fewer than 10M parameters, significantly less than their Transformer-like counterparts. However, it is also clear from the figure that the accuracy of graph-based models tends to lag behind that of Transformer models. Fig. 2 (a) shows that the minimal features needed to describe the 3D structure include bonds, angles, and torsions. All QC methods requires 3D structural information [17], which directly influence electronic distribution [18]. Unfortunately, Fig. 2 (b) shows the traditional graph convolutional neural networks focus solely on 1-hop information, making it challenging to predict molecular properties accurately.

**Fig. 2.**
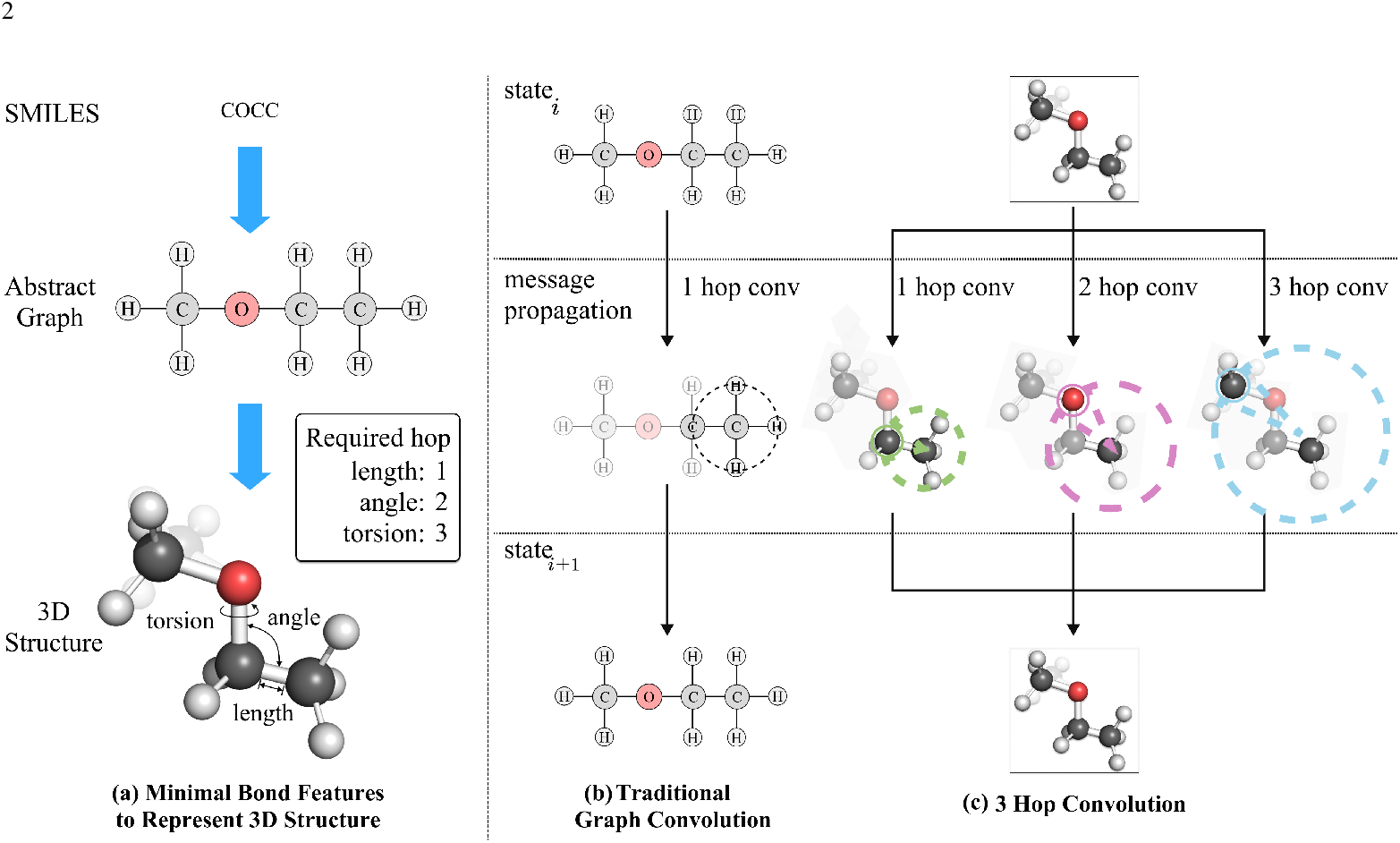
To accurately represent 3D structures, it is necessary to incorporate at least 3-hop features, which include bond length, bond angle, and torsion angle. This means that traditional Graph Convolutional Networks (GCNs) that only utilize 1-hop features fall short in representing 3D structures. MetaGIN, on the other hand, efficiently uses 3-hop features, satisfying the bare minimum requirements for 3D structure representation.

Moreover, as molecular databases such as ZINC [19] and ChemSpider [20] continue to expand, now containing hundreds of millions of small molecules, an increasing number of these molecules are missing annotated 3D structural information. This absence represents a significant challenge in the realm of molecular property prediction, particularly given the wealth of information that 3D structures provide for understanding molecular properties. This challenge highlights an immediate need for the development of models capable of operating effectively without the dependence on 3D structural data. Such models are particularly crucial for training and inference on a substantial subset of molecules where this structural information is not readily available.

Inspired by the foregoing discussion, we present MetaGIN, a lightweight framework designed for molecular property prediction without using 3D structure. By incorporating graph convolution and a virtual node block into the MetaFormer architecture [21], MetaGIN not only reduces the number of parameters but also achieves performance comparable to the Transformer-like models. Additionally, as Fig. 2 (c) shows, we propose a 3-hop convolution mechanism to capture essential molecular features such as bond, angle, and torsion, effectively describing the 3D structure of molecules. The implementation of 3-hop convolution contributes to improved performance over traditional graph convolutional neural networks.

## 2. Method

### 2.1 Overview of MetaGIN

As depicted in Fig. 3 (a), the MetaGIN architecture processes molecular inputs to yield their respective molecule properties. MetaGIN is structured over *L* layers, each layer takes two embedding as input: 1) the graph embedding *G*^(*l*)^ and 2) the node embedding *X*^(*l*)^. The graph embedding *G*^(*l*)^ ∈ *R*^*d*^ is initialized as a zero vector to represent the graph state. And the initial node embedding *X*^(0)^ ∈ *R*^*n*×*d*^ is generated by the open graph benchmark (OGB) toolkit [9, 22] firstly and then passed through an embedding layer to obtain the proper representation of node features. The node embedding *X*^(*l*)^ and graph embedding *G*^(*l*)^ are updated and passed to the next layer as input in each layer.

**Fig. 3.**
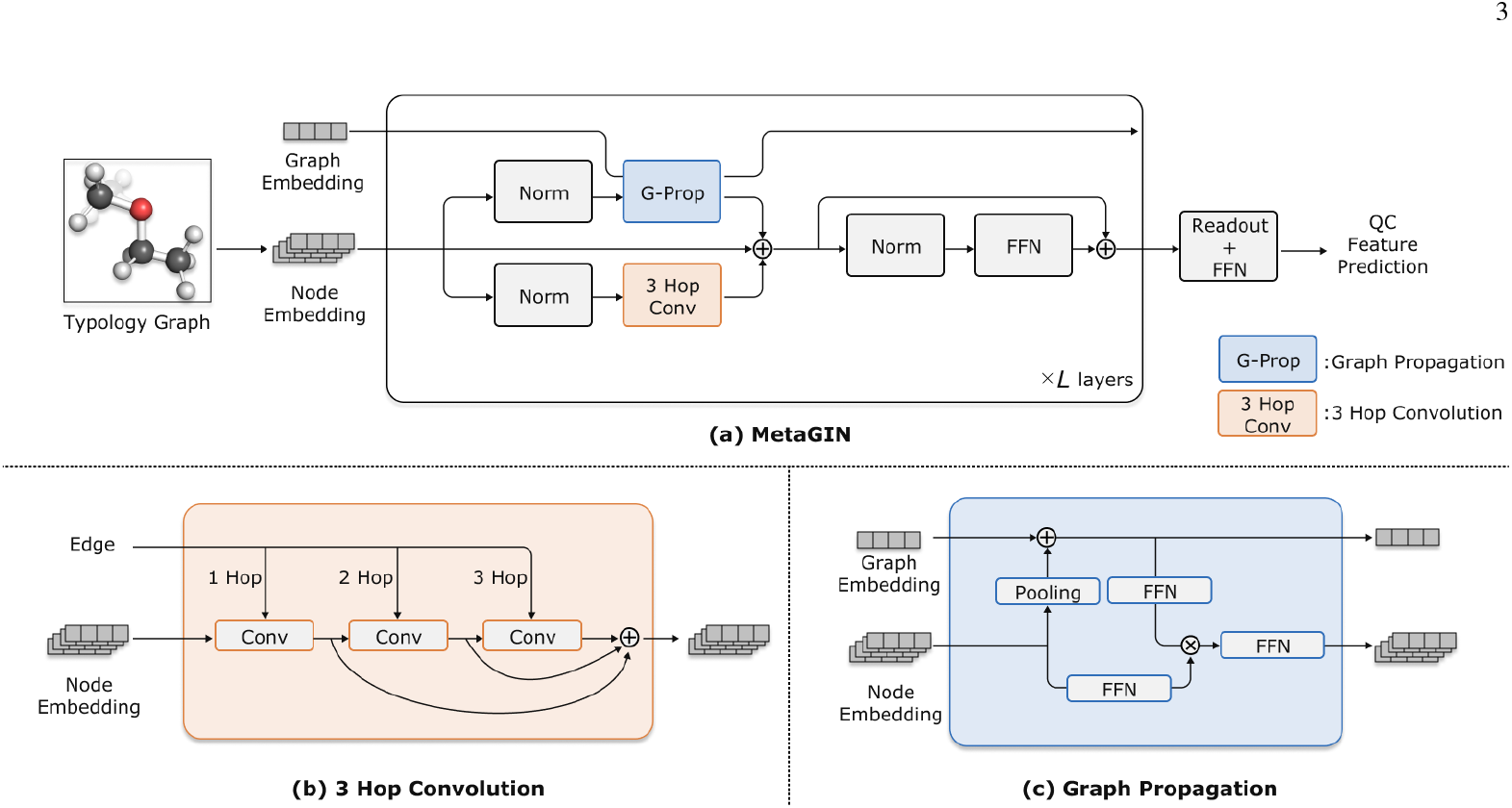
(a) The architecture of MetaGIN is based on MetaFormer, which includes both a token mixer block and a feed-forward neural (FFN) network block. Particularly, MetaGIN introduces two types of token mixer blocks: (b) the 3-hop convolution block and the (c) graph propagation block.

Drawing inspiration from MetaFormer [21], MetaGIN incorporates the graph neural network related methods. These methods serve as the token mixture block, thereby bringing the core architecture principles of transformers into the realm of graph neural networks. Within the graph token mixture, we introduce two variants: 3 hop convolution block and graph propagation block. These distinct block types will be detailed in Section 2.2 and Section 2.3.

### 2.2 3 Hop Convolution

The 3 Hop Convolution block is responsible for the local information propagation. As shown in Fig. 3 (b), 3 hop convolution is applied to obtain the ability to learn the 3D structure of molecules. The 3 hop convolution block takes the node embedding *X*^(*l*)^ and the edge information *E* as input. There are three types of edge information: 1) 1 hop edge, 2) 2 hop edge, and 3) 3 hop edge. The 1 hop edge can be obtained by OGB toolkit [9,22] directly which is 5 type bond features like bond type and chirality. Different from the 1 hop feature, the 2 hop and 3 hop edge features both are 1 dimensional features that are preprocessed to indicate the number of available paths between two nodes. All the hop edge features *E* = {*E*_1_, *E*_2_, *E*_3_} are embedding to the same dimention as the node embedding *X*^(*l*)^.

To enhance the 3D structural description of molecules, we employ three types of edge features within our 3 hop convolution blocks. These convolution blocks are strategically linked in sequence to inherit information from previous hops. The node embedding *X*^(*l*)^ undergoes an update to *X*^(*l*+1)^ through the three-hop convolution process, which can be described as follows:

First, we perform the three continuous local convolutions:

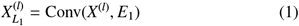

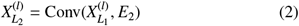

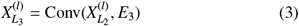

Here, 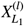, 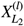, and 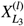 represent the node embeddings updated by one-hop, two-hop, and three-hop convolutions, respectively.

Next, these results are combined to update the node embedding for the next layer, 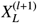, with a strong emphasis on the connection between them:

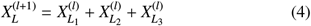

In these equations, *Conv* denotes the graph convolution function, here we choose GIN [16] as the graph convolution function, and for each node *v* ∈ *V* the propagation process can be described as follows:

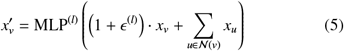

In these equations, *V* is the set of nodes, N(*v*) represents the neighbors of node *v*, and ϵ is a learnable parameter. The result, 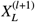, embodies the node embedding updated by the local convolution block.

### 2.3 Graph Propagation

While the 3 hop convolution excels in capturing localized information within molecules, it falls short in perceiving the global context of the entire molecule. To address this limitation and expedite message propagation across the molecular structure, we introduce the concept of global propagation. This approach draws inspiration from the virtual node module [16], enabling a comprehensive understanding of the entire molecule. Instead of relying on simplistic broadcast mechanisms, we employ gating mechanisms to control the graph information received by each node. The following content will provide a detailed exposition of our global convolution methodology.

The core idea behind the virtual node is to learn a graph representation and subsequently broadcast this updated representation to each node within the graph. However, a limitation of this approach is that each node receives identical information during the broadcast, leaving no room for differential reception based on the node’s inherent properties or state. To overcome this challenge, we apply the gating mechanisms within the broadcast of the graph representation, enabling nodes to control the information they receive from the graph embedding through their node embedding. This mechanism not only facilitates the exchange of global information but also allows different nodes to exhibit varying degrees of sensitivity to the information, maintaining the uniqueness in the perception of each node within the global context.

To amplify the capacity of the model to comprehend the global context of molecules, the graph propagation block engages two inputs, specifically the node embedding *X*^(*l*)^ and the graph embedding *G*^(*l*)^.

Initially, a pooling operation, Pool, is performed on the node embedding *X*^(*l*)^, to obtain a new graph embedding of the current layer. And subsequently, the newly obtained graph embedding is subjected to an update mechanism to form the updated representative vector 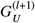. Due to the sum pooling being more representative than the average pooling and max pooling [16], we choose the sum pooling as the pooling operation.

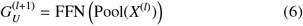

This updated representative vector 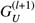 is then disseminated to each node. Here, each node’s embedding acts as the gate 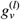 controlling the received information from 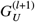 based on its specific properties. the gate *g*^(*l*)^ is optimized by their node embedding *X*^(*l*)^. The gate *g*^(*l*)^ and the propagated node embedding *X*^(*l*+1)^ are updated as follows:

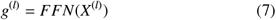

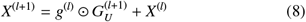

Here, the ⊙ represents element-wise multiplication, and FFN denotes the feedforward network.

After the dissemination and modulation processes, each node embedding undergoes refinement through a feedforward network, FFN, facilitating the comprehensive integration of refined global information in the subsequent layers of the model.

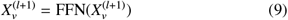

This mechanism not only enables effective global information exchange but also maintains the distinctiveness in the information reception of each node within the global context, enriching the model’s understanding of the overall molecular structure.

This structured depiction of the Global Convolution aims to delineate the interrelation between its constituents and their sequential transformation, offering clear insight into the inherent workflow of the methodology.

## 3 Results

We evaluate the performance and robustness of MetaGIN on two key benchmarks: the PCQM4Mv2 dataset [9, 22] and the MoleculeNet benchmark [23]. We then conduct an ablation study to identify the crucial components of MetaGIN. Additionally, we examine the insensitivity of MetaGIN to chiral data, highlighting its robustness and guiding future research. The following subsections provide detailed insights into each of these aspects.

### 3.1 Performance on PCQM4Mv2

The PCQM4Mv2 dataset [9,22], comprising over 337 million molecules, aids in ML-driven predictions of DFT-calculated HOMO-LUMO energy gaps using 2D molecular graphs. We chose this dataset for evaluation due to its widespread recognition and the continuous efforts to advance models based on it, making results on this dataset more convincing and reflective of a model’s practical robustness.

In Table 1, we categorize the evaluated algorithms into two groups: graph based models and Transformer-like architectures, focusing solely on models that do not utilize 3D structural as described in Section 1. The primary metric for evaluation is the Mean Absolute Error (MAE), where lower values indicate superior performance.

**Table 1.**
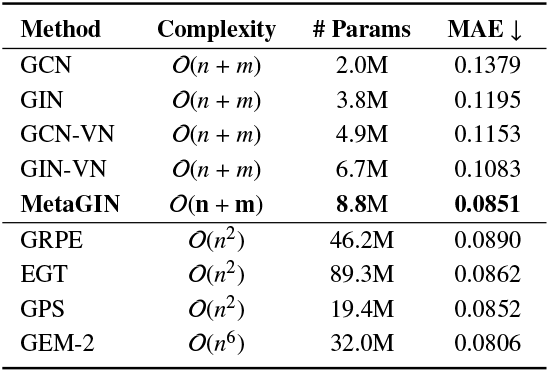
HOMO/LUMO gap prediction on the PCQM4Mv2 dataset.

| Method | Complexity | # Params | MAE ↓ |
| --- | --- | --- | --- |
| GCN | $O(n + m)$ | 2.0M | 0.1379 |
| GIN | $O(n + m)$ | 3.8M | 0.1195 |
| GCN-VN | $O(n + m)$ | 4.9M | 0.1153 |
| GIN-VN | $O(n + m)$ | 6.7M | 0.1083 |
| **MetaGIN** | **$O(n + m)$** | **8.8M** | **0.0851** |
| GRPE | $O(n^2)$ | 46.2M | 0.0890 |
| EGT | $O(n^2)$ | 89.3M | 0.0862 |
| GPS | $O(n^2)$ | 19.4M | 0.0852 |
| GEM-2 | $O(n^6)$ | 32.0M | 0.0806 |

Among the models that operate without 3D information, MetaGIN achieves a MAE of 0.0851. This surpasses other models like GCN, GIN, GCN-VN, and GIN-VN, all of which operate under the same computational complexity of O(*n* + *m*) but yield higher MAE values ranging from 0.1083 to 0.1379. Furthermore, MetaGIN is parameter efficient, requiring only 8.8M parameters, in contrast to models like GRPE, EGT, and GPS, which require between 46.2M and 89.3M parameters.

Interestingly, the performance of MetaGIN is also competitive with models having higher computational complexity like GRPE, EGT, and GPS that operate in O(*n*^2^), and GEM-2, which operates in O(*n*^6^). The Mean Absolute Error (MAE) of MetaGIN closely approaches that of the optimal model GEM-2, ranking second in the table. This highlights the model’s effective balance between computational efficiency and performance.

In summary, the empirical results on the PCQM4Mv2 dataset validate the robustness and efficiency of MetaGIN, positioning it as a compelling choice for tasks that demand both high performance and computational economy.

### 3.2 Performance on MoleculeNet

Although Section 3.1 has already demonstrated good performance of our model on the PCQM4Mv2 dataset, to further validate the generalizability of MetaGIN across different datasets, we extend our evaluation to the MoleculeNet benchmark [23].

Table 2 and Table 3 provides a comprehensive evaluation of various machine learning models on MoleculeNet benchmark, focusing on both classification and regression tasks. All the dataset is sorted by the number of molecules in the dataset. The table is divided into two primary sections, delineated by a horizontal line. The upper section lists models that have not been pre-trained, including ECFP [24], TF Robust [25], GraphConv [26], Weave [27], SchNet [28], MGCN [29], AttentiveFP [30], and TrimNet [31]. While the lower section comprises models that have undergone pre-training. Performance metrics for each model are displayed in their respective cells, including MPNN [32], DMPNN [33], FunQG-MPNN [34], FunQG-DMPNN [34], and MetaGIN. The best-performing model in the pre-trained section is highlighted in bold, while in the nonpre-trained section, models are color-coded. Specifically, models outperforming their pre-trained counterparts are marked in green, while the others are indicated in red.

**Table 2.** Regression tasks on MoleculeNet datasets.

|  | FreeSolv | ESOL | Lipophilicity |
| --- | --- | --- | --- |
| # Mols | 642 | 1128 | 4200 |
| **Without pretraining** |  |  |  |
| ECFP | 5.275(0.751) | 2.359(0.454) | 1.188(0.061) |
| TF_Robust | 4.122(0.085) | 1.722(0.038) | 0.909(0.060) |
| GraphConv | 2.900(0.135) | 1.068(0.050) | 0.712(0.049) |
| Weave | 2.398(0.250) | 1.158(0.055) | 0.813(0.042) |
| SchNet | 3.215(0.755) | 1.045(0.064) | 0.909(0.098) |
| MGCN | 3.349(0.097) | 1.266(0.147) | 1.113(0.041) |
| AttentiveFP | 2.030(0.420) | 0.853(0.060) | 0.650(0.030) |
| TrimNet | 2.529(0.111) | 1.282(0.029) | 0.702(0.008) |
| **Pretrained** |  |  |  |
| MPNN | 2.185(0.952) | 1.167(0.430) | 0.885(0.030) |
| DMPNN | 2.177(0.914) | 0.980(0.258) | 0.653(0.046) |
| FunQG-MPNN | 1.542(0.460) | 0.879(0.091) | 0.638(0.020) |
| FunQG-DMPNN | 1.501(0.376) | 0.818(0.047) | 0.622(0.028) |
| **MetaGIN** | **1.397(0.062)** | **0.780(0.061)** | **0.532(0.013)** |

**Table 3.**
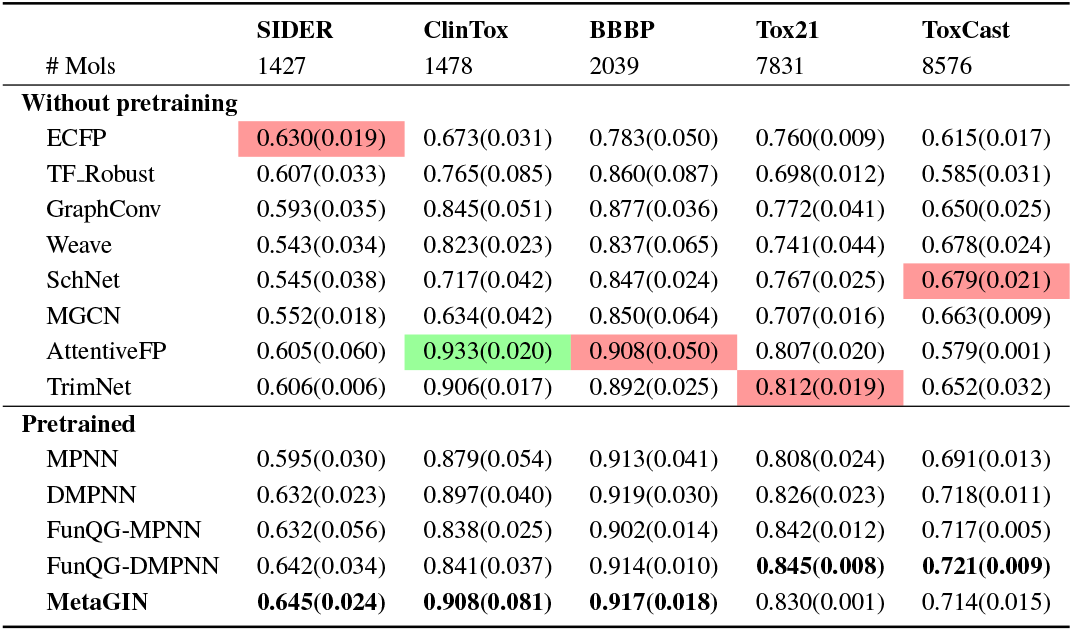
Classification tasks on MoleculeNet datasets.

| # Mols | SIDER<br>1427 | ClinTox<br>1478 | BBBP<br>2039 | Tox21<br>7831 | ToxCast<br>8576 |
| --- | --- | --- | --- | --- | --- |
| **Without pretraining** |  |  |  |  |  |
| ECFP | 0.630(0.019) | 0.673(0.031) | 0.783(0.050) | 0.760(0.009) | 0.615(0.017) |
| TF_Robust | 0.607(0.033) | 0.765(0.085) | 0.860(0.087) | 0.698(0.012) | 0.585(0.031) |
| GraphConv | 0.593(0.035) | 0.845(0.051) | 0.877(0.036) | 0.772(0.041) | 0.650(0.025) |
| Weave | 0.543(0.034) | 0.823(0.023) | 0.837(0.065) | 0.741(0.044) | 0.678(0.024) |
| SchNet | 0.545(0.038) | 0.717(0.042) | 0.847(0.024) | 0.767(0.025) | 0.679(0.021) |
| MGCN | 0.552(0.018) | 0.634(0.042) | 0.850(0.064) | 0.707(0.016) | 0.663(0.009) |
| AttentiveFP | 0.605(0.060) | 0.933(0.020) | 0.908(0.050) | 0.807(0.020) | 0.579(0.001) |
| TrimNet | 0.606(0.006) | 0.906(0.017) | 0.892(0.025) | 0.812(0.019) | 0.652(0.032) |
| **Pretrained** |  |  |  |  |  |
| MPNN | 0.595(0.030) | 0.879(0.054) | 0.913(0.041) | 0.808(0.024) | 0.691(0.013) |
| DMPNN | 0.632(0.023) | 0.897(0.040) | 0.919(0.030) | 0.826(0.023) | 0.718(0.011) |
| FunQG-MPNN | 0.632(0.056) | 0.838(0.025) | 0.902(0.014) | 0.842(0.012) | 0.717(0.005) |
| FunQG-DMPNN | 0.642(0.034) | 0.841(0.037) | 0.914(0.010) | **0.845(0.008)** | **0.721(0.009)** |
| **MetaGIN** | **0.645(0.024)** | **0.908(0.081)** | **0.917(0.018)** | 0.830(0.001) | 0.714(0.015) |

Upon examining the table, it is clear that our model, denoted as MetaGIN, significantly outperforms other architectures across the majority of datasets in both classification and regression tasks. In situations where MetaGIN does not secure the top position, its performance remains competitive, closely trailing the best-performing models. For instance, in the ToxCast dataset for classification tasks, our model’s performance is nearly identical to the leading model, FunQG-DMPNN, with a negligible difference of only 0.007. This gap falls within the variance of the leading model, indicating that MetaGIN may potentially outperform FunQG-DMPNN in certain cases. Likewise, in the ClinTox dataset, MetaGIN is only 0.025 behind the leading model, AttentiveFP [30]. This difference is within the variance of our model, underscoring its competitive edge. These findings affirm the model’s versatility and effectiveness, particularly when it is pre-trained on the PCQM4Mv2 dataset.

Several key insights emerge from this comprehensive evaluation. Firstly, the influence of pre-training is evident, particularly in the superior performance of our model, MetaGIN. The model not only excels in various tasks but also consistently ranks among the top performers, thereby underscoring the effectiveness of pre-training when combined with our architectural choices.

Secondly, the robustness of MetaGIN is noteworthy, as it manages to deliver high performance across multiple datasets and tasks. This points to the model’s capacity for generalization, making it a reliable choice for both classification and regression tasks.

A notable observation is the particularly strong performance of MetaGIN in regression tasks. The model’s greater edge in these tasks could be attributed to the nature of the pre-training dataset,

PCQM4Mv2, suggesting that the type of tasks in the pre-training dataset may influence the model’s effectiveness. This is an aspect that warrants further investigation in future work.

The evaluation clearly signifies that MetaGIN sets a new benchmark in the field, particularly when pre-trained on the PCQM4Mv2 dataset.

### 3.3 Ablation Studies

After conducting extensive ablation studies to evaluate the different components of MetaGIN, we present our findings in Table 4. Each row in the table represents a unique model configuration, detailing the number of hops, repetition rate of convolutions, depth (also represented as “Depth” in the table), width (also represented as “Width” in the table), number of parameters (#Params), and Validation Mean Absolute Error (MAE) scores. It should be noted that a lower MAE is preferable.

**Table 4.** Ablation studies.

| Hop | Repeat | Depth | Width | #Params | MAE ↓ |
| --- | --- | --- | --- | --- | --- |
| 1 | 1 | 4 | 256 | 2.82M | 0.0920 |
| 2 | 1 | 4 | 256 | 3.63M | 0.0881 |
| 3 | 1 | 4 | 256 | 4.45M | 0.0871 |
| 1 | 3 | 4 | 256 | 4.29M | 0.0883 |
| 3 | 1 | 8 | 256 | 8.92M | 0.0855 |
| 3 | 1 | 4 | 512 | 8.87M | 0.0851 |

Our observations are grouped into two primary categories. First, we evaluated the impact of varying the number of hops and the repetition rate of convolutions. Comparisons were made between configurations with 1-hop, 2-hop, and 3-hop convolutions, as well as a configuration where a 1-hop convolution was repeated three times. Increasing the number of hops led to improved performance, thereby emphasizing the importance of multi-hop graph convolutions in our framework. Conversely, simply repeating a 1-hop convolution three times did not yield a significant performance improvement, as reflected by its Validation MAE score of 0.0883.

In the second category, we examined the trade-offs between model depth and width while keeping the total number of parameters constant. Two particular configurations were of interest: one with a depth of 8 layers and another with a width of 512 units. Both configurations showed comparable performance, with the latter having a slight edge as evidenced by its Validation MAE score of 0.0851. This suggests that increasing the model’s width may offer some performance benefits when the total parameter count remains constant.

The results of these ablation experiments affirm the robust performance of MetaGIN and provide valuable insights into the relative contributions of its constituent elements.

### 3.4 Discussions on Chirality

We initiated an experiment to better understand how molecular chirality impacts the performance of our model. The findings are presented in a two-part figure, as shown in Fig. 4. The first part, Fig. 4 (a), highlights two chiral configurations of a small molecule. Despite a minimal notational difference in their SMILES representations—namely ‘@’ versus ‘@@’— the 3D spatial configurations vary significantly. For instance, the distance between the red oxygen and blue nitrogen atoms is 5.9 Å in one configuration and 5.1 Å in the other.

**Fig. 4.**
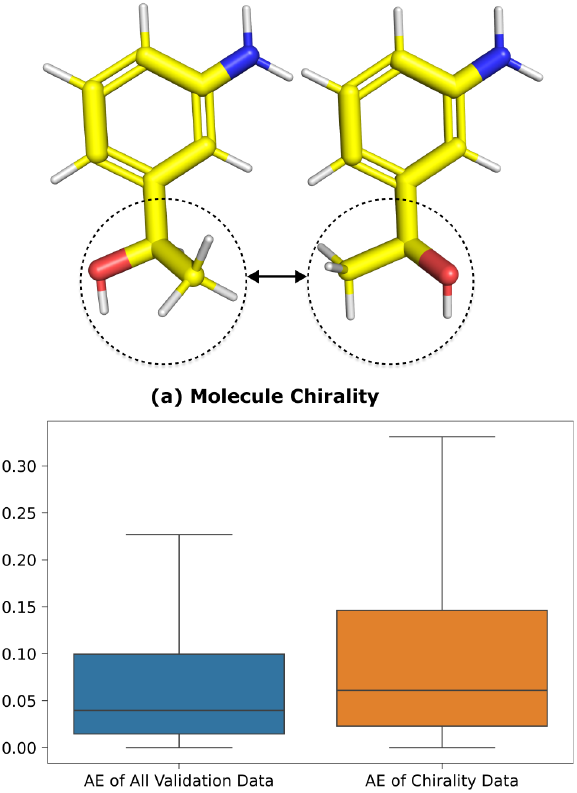
(a) Chirality is especially important in the field of biochemistry because the biological properties of chiral molecules can be drastically different between their enantiomers. (b) Indeed, the presence of chirality leads to a decrease in accuracy when predicting the HOMO-LUMO gap.

Turning to Fig. 4 (b), the boxplot visualizes the Absolute Error (AE) distribution for the general validation set alongside a subset composed of chiral molecules from the PCQM4Mv2 dataset. Unfortunately, our model’s performance is less robust when dealing with chiral molecules, as evidenced by both the median AE and the distribution trend.

To sum up, the experimental findings spotlight the limitations of our model in accurately predicting chiral molecules, possibly due to the absence of 3D structural data. We’re considering the incorporation of such structural data in future versions of the model to improve its sensitivity to chirality.

## 4 Conclusion

In this work, we present MetaGIN, a lightweight framework for molecular property prediction. In PCQM4Mv2 dataset, MetaGIN not only matches the performance of Transformer-like models but also maintains superior efficiency in both parameter count and computational time. For the majority of datasets in the MoleculeNet benchmark, MetaGIN achieves state-of-the-art performance, to the best of our knowledge. Additionally, inspired by the characteristics of quantum chemistry (QC) methods, we introduce 3-hop convolution to more accurately describe the 3D structure of molecules. Our ablation studies show the superiority of 3-hop convolution when compared to traditional graph convolutional neural networks that only consider 1-hop information. The success of the 3-hop convolution indicates that domain knowledge can assist in the design of more effective models.

Furthermore, our investigation into molecular chirality has revealed certain limitations of our model. Specifically, MetaGIN demonstrates a diminished ability to accurately predict properties of chiral molecules, as evidenced in our experiments outlined in Fig. 4. These findings point to potential areas for future research, including the incorporation of 3D structural data to enhance the model’s sensitivity to chirality.

In summary, MetaGIN represents a significant advancement in the field of molecular property prediction, effectively marrying high performance with computational efficiency. Our model not only establishes new benchmarks across multiple datasets but also introduces architectural innovations that have the potential to guide future research in this domain.

## ACKNOWLEDGMENT

This research has been facilitated by the National Key R&D Program of China under the grants No. 2019YFA0905700 and 2021YFC2101500, as well as a grant from the National Science Foundation of China, grant number 62072283.

## References

1. Lin X, Li X, Lin X. A Review on Applications of Computational Methods in Drug Screening and Design. Molecules, 2020, 25(6): 1375

2. Hann M M, Leach A R, Harper G. Molecular Complexity and Its Impact on the Probability of Finding Leads for Drug Discovery. Journal of Chemical Information and Computer Sciences, 2001, 41(3): 856–864

3. Manallack D T, Prankerd R J, Yuriev E, Oprea T I, Chalmers D K. The significance of acid/base properties in drug discovery. Chemical Society Reviews, 2013, 42(2): 485–496

4. Conceptual Density Functional Theory — Chemical Reviews. https://pubs.acs.org/doi/full/10.1021/cr990029p

5. Motta M, Zhang S. Ab initio computations of molecular systems by the auxiliary-field quantum Monte Carlo method. WIREs Computational Molecular Science, 2018, 8(5): e1364

6. Kümmel H G. A biography of the coupled cluster method. International Journal of Modern Physics B, 2003, 17(28): 5311–5325

7. Zhang X, Chen C, Meng Z, Yang Z, Jiang H, Cui X. CoAtGIN: Marrying Convolution and Attention for Graph-based Molecule Property Prediction. In: 2022 IEEE International Conference on Bioinformatics and Biomedicine (BIBM). December 2022, 374–379

8. Wang Z, Wang Y, Zhang X, Meng Z, Yang Z, Zhao W, Cui X. Graphbased Reaction Classification by Contrasting between Precursors and Products. In: 2022 IEEE International Conference on Bioinformatics and Biomedicine (BIBM). December 2022, 354–359

9. Hu W, Fey M, Ren H, Nakata M, Dong Y, Leskovec J. OGB-LSC: A Large-Scale Challenge for Machine Learning on Graphs, October 2021

10. Vaswani A, Shazeer N, Parmar N, Uszkoreit J, Jones L, Gomez A N, Kaiser Ł, Polosukhin I. Attention is all you need. In: Guyon I, Luxburg U V, Bengio S, Wallach H, Fergus R, Vishwanathan S, Garnett R, eds, Advances in Neural Information Processing Systems. 2017

11. Liu L, He D, Fang X, Zhang S, Wang F, He J, Wu H. GEM-2: Next Generation Molecular Property Prediction Network by Modeling Fullrange Many-body Interactions, October 2022

12. Dwivedi V P, Luu A T, Laurent T, Bengio Y, Bresson X. Graph Neural Networks with Learnable Structural and Positional Representations. In: International Conference on Learning Representations. October 2021

13. Hussain M S, Zaki M J, Subramanian D. Global Self-Attention as a Replacement for Graph Convolution. In: Proceedings of the 28th ACM SIGKDD Conference on Knowledge Discovery and Data Mining. August 2022, 655–665

14. Park W, Chang W G, Lee D, Kim J, Hwang S W. GRPE: Relative Positional Encoding for Graph Transformer. In: ICLR2022 Machine Learning for Drug Discovery. April 2022

15. Kipf T N, Welling M. Semi-Supervised Classification with Graph Convolutional Networks, February 2017

16. Xu* K, Hu* W, Leskovec J, Jegelka S. How Powerful are Graph Neural Networks? In: International Conference on Learning Representations. September 2018

17. Thiel W. Semiempirical quantum–chemical methods. WIREs Computational Molecular Science, 2014, 4(2): 145–157

18. Bannwarth C, Caldeweyher E, Ehlert S, Hansen A, Pracht P, Seibert J, Spicher S, Grimme S. Extended tight-binding quantum chemistry methods. Wiley Interdisciplinary Reviews: Computational Molecular Science, 2021, 11(2): e1493

19. Irwin J J, Tang K G, Young J, Dandarchuluun C, Wong B R, Khurelbaatar M, Moroz Y S, Mayfield J, Sayle R A. ZINC20—A Free Ultralarge-Scale Chemical Database for Ligand Discovery. Journal of Chemical Information and Modeling, 2020, 60(12): 6065–6073

20. Pence H E, Williams A. ChemSpider: An Online Chemical Information Resource. Journal of Chemical Education, 2010, 87(11): 1123–1124

21. Yu W, Luo M, Zhou P, Si C, Zhou Y, Wang X, Feng J, Yan S. MetaFormer Is Actually What You Need for Vision. In: Proceedings of the IEEE/CVF Conference on Computer Vision and Pattern Recognition. 2022, 10819–10829

22. Hu W, Fey M, Zitnik M, Dong Y, Ren H, Liu B, Catasta M, Leskovec J. Open graph benchmark: Datasets for machine learning on graphs. In: Larochelle H, Ranzato M, Hadsell R, Balcan M, Lin H, eds, Advances in Neural Information Processing Systems. 2020, 22118–22133

23. Wu Z, Ramsundar B, Feinberg E N, Gomes J, Geniesse C, Pappu A S, Leswing K, Pande V. MoleculeNet: A benchmark for molecular machine learning †Electronic supplementary information (ESI) available. See DOI: 10.1039/c7sc02664a. Chemical Science, 2017, 9(2): 513–530

24. Ramsundar B, Kearnes S, Riley P F, Webster D, Konerding D, Pande V. Massively Multitask Networks for Drug Discovery. ArXiv, 2015

25. Rogers D, Hahn M. Extended-Connectivity Fingerprints. Journal of Chemical Information and Modeling, 2010, 50(5): 742–754

26. Kipf T N, Welling M. Semi-Supervised Classification with Graph Convolutional Networks. In: International Conference on Learning Representations. November 2016

27. Kearnes S, McCloskey K, Berndl M, Pande V, Riley P. Molecular graph convolutions: Moving beyond fingerprints. Journal of Computer-Aided Molecular Design, 2016, 30(8): 595–608

28. Schütt K, Kindermans P J, Sauceda Felix H E, Chmiela S, Tkatchenko A, Müller K R. SchNet: A continuous-filter convolutional neural network for modeling quantum interactions. In: Advances in Neural Information Processing Systems. 2017

29. Lu C, Liu Q, Wang C, Huang Z, Lin P, He L. Molecular property prediction: A multilevel quantum interactions modeling perspective. In: Proceedings of the Thirty-Third AAAI Conference on Artificial Intelligence and Thirty-First Innovative Applications of Artificial Intelligence Conference and Ninth AAAI Symposium on Educational Advances in Artificial Intelligence, AAAI’19/IAAI’19/EAAI’19. January 2019, 1052–1060

30. Xiong Z, Wang D, Liu X, Zhong F, Wan X, Li X, Li Z, Luo X, Chen K, Jiang H, Zheng M. Pushing the Boundaries of Molecular Representation for Drug Discovery with the Graph Attention Mechanism. Journal of Medicinal Chemistry, 2020, 63(16): 8749–8760

31. Liaw R, Liang E, Nishihara R, Moritz P, Gonzalez J E, Stoica I. Tune: A Research Platform for Distributed Model Selection and Training, July 2018

32. Gilmer J, Schoenholz S S, Riley P F, Vinyals O, Dahl G E. Neural Message Passing for Quantum Chemistry. In: Proceedings of the 34th International Conference on Machine Learning. July 2017, 1263–1272

33. Yang K, Swanson K, Jin W, Coley C, Eiden P, Gao H, Guzman-Perez A, Hopper T, Kelley B, Mathea M, Palmer A, Settels V, Jaakkola T, Jensen K, Barzilay R. Analyzing Learned Molecular Representations for Property Prediction. Journal of Chemical Information and Modeling, 2019, 59(8): 3370–3388

34. Hajiabolhassan H, Taheri Z, Hojatnia A, Yeganeh Y T. FunQG: Molecular Representation Learning via Quotient Graphs. Journal of Chemical Information and Modeling, 2023, 63(11): 3275–3287

